# Ribosomal DNA methylation as stable biomarkers for detection of cancer in plasma

**DOI:** 10.1101/651497

**Authors:** Xianglin Zhang, Huan Fang, Wei Zhang, Bixi Zhong, Yanda Li, Xiaowo Wang

**Affiliations:** Department of Automation, Center for Synthetic and Systems Biology, Tsinghua University; Ministry of Education Key Laboratory of Bioinformatics; Bioinformatics Division, BNRIST, Beijing 100084, China

**Keywords:** ribosomal DNA, liquid biopsy, biomarker, cancer detection, DNA methylation, cfDNA, liver cancer, colon cancer, lung cancer

## Abstract

**Background:** Recently, liquid biopsy for cancer detection has pursued great progress. However, there are still a lack of high quality markers. It is a challenge to detect cancer stably and accurately in plasma cell free DNA (cfDNA), when the ratio of cancer signal is low. Repetitive genes or elements may improve the robustness of signals. In this study, we focused on ribosomal DNA which repeats hundreds of times in human diploid genome and investigated performances for cancer detection in plasma.

**Results:** We collected bisulfite sequencing samples including normal tissues and 4 cancer types and found that intergenic spacer (IGS) of rDNA has high methylation levels and low variation in normal tissues and plasma. Strikingly, IGS of rDNA shows significant hypo-methylation in tumors compared with normal tissues. Further, we collected plasma bisulfite sequencing data from 224 healthy subjects and cancer patients. Means of AUC in testing set were 0.96 (liver cancer), 0.94 (lung cancer and), 0.92 (colon cancer) with classifiers using only 10 CpG sites. Due to the feature of high copy number, when liver cancer plasma WGBS was down-sampled to 10 million raw reads (0.25× whole genome coverage), the prediction performance decreased only a bit (mean AUC=0.93). Finally, methylation of rDNA could also be used for monitor cancer progression and treatment.

**Conclusion:** Taken together, we provided the high-resolution map of rDNA methylation in tumors and supported that methylation of rDNA was a competitive and robust marker for detecting cancer and monitoring cancer progression in plasma.

## Background

Nowadays, liquid biopsy for cancer managements including cancer detection, monitoring and treatment selection develops quickly due to the improvement of genomic and molecular methods, as well as its advantages of non-invasion and overcoming the limitation of tumor heterogeneity compared with traditional tissue biopsy [1–3]. In liquid biopsy, cell-free DNA (cfDNA) becomes one of the most widely studied analysts because of its stability, abundance and potential applications in precision oncology[2]. cfDNA can be released from cells into body fluids through apoptosis, necrosis and active secretion[2, 3]. In the same way, circulating tumor DNA (ctDNA) is released from malignant tumors during the progress of cancer. Identifying ctDNA from other cfDNA with tumor DNA-specific features is the principle of cancer detection. Previous studies showed that point mutations, copy number alterations, structural variants, methylation changes, nucleosomes positioning changes and fragment length differences reflected by tumor DNA can be used as diagnostic markers[2]. Among these markers, aberrant DNA methylation provides indispensable information for cancer managements and have been widely studied [2, 4–6].

Bisulfite conversion sequencing is considered as “gold standard” of methylation sequencing[6] and have been widely used to investigate DNA methylation patterns in single base-pair resolution[7]. Whole genome bisulfite sequencing (WGBS) and reduced representation bisulfite sequencing (RRBS) are two common bisulfite conversion-based technologies. WGBS is fairly accurate, reproducible to obtain methylation profiles of whole genome, but is costly and low coverage[7]. RRBS is relatively cost-effective, but covers only 10% of whole genome and is GC-biased[7]. They have been used for cfDNA researches in liver cancer, lung cancer and colon cancer[8, 9].

Finding accurate DNA methylation markers is a key for cancer detection [10–12]. When there is low tumor burden in plasma, bisulfite sequencing of cfDNA from several milliliter plasma might produce unstable results [2]. However, there are some repeat units with high copy number in human genome, and selecting them as biomarkers may enhance the robustness of signals and improve cancer detection performances. In this study, we paid attention to human ribosomal DNA (rDNA) repeat unit, which repeats hundreds of times in diploid human genome.

Human rDNA clusters on the short arms of the five acrocentric chromosomes (chromosomes 13, 14, 15, 21, 22), and contains a 13-kb transcribed region and a 30-kb intergenic spacer (IGS). Transcribed region encodes 18S, 5.8S and 28S ribosomal RNA. IGS contains multiple mobile elements, transcription factor binding sites like TP53 binding sites and sequences similar to scaffold attachment regions, which indicates IGS may play a role in enhancement of transcription, recombination and chromosome organization[13]. Recently, a study identified 49 conserved regions in IGS through comparative genome in multiple species[14] and revealed the regulation potentials of IGS.

rDNA is the fundamental genomic element to form nucleolus where ribosomes assemble together[15, 16] and its transcriptional RNA product - ribosomal RNA is indispensable parts of ribosome, which suggests rDNA stability is vital for normal biology processes. Copy number variation of rDNA balances rDNA dosage [17]and is associated with the expression of several functional coherent gene sets[18]. rDNA copy number amplification or deletion is related to tumor genetic context, nucleolus activity, proliferation and protein activity in different cancer types[19–22]. The expression of rDNA also shows aberrant patterns in prostate cancer, cervical cancer, breast cancer and myelodysplastic syndromes[23–26]. Overexpression of pre-45S ribosomal RNA promotes colon cancer and is associated with poor survival of colorectal patients[27].

Epigenetic regulation is an important mechanism of keeping rDNA stable [28–30] and its disruption is highly related with aging and cancer[30–33]. Researchers found that transcribed region of rDNA is un-methylated, while IGS is highly methylated, and relatively sharp boundaries are formed between two regions in blood cells[34]. In contrast, IGS becomes partially de-methylated in lung cancer[35] and the promoters or transcribed domain of rDNA show hyper-methylation in tumor tissues of breast cancer, ovarian cancer, endometrial carcinoma and colorectal cancer [36–40]. Hyper-methylation in rDNA locus predict longer survival in ovarian cancer, endometrial carcinoma[38, 39]. In addition, histone modification analysis shows that, liver cancer cells have distinct epigenetic pattern compared with normal liver cells[41]. Previous study has integrated ChIP-seq, DNase-seq, MNase-seq and RNA-seq data to study the epigenetic and transcriptional patterns of rDNA in several cell lines[42], however, there is no methylation map of rDNA repeat unit including both transcribed region and IGS in single CpG resolution, and there is no study focusing on the potential usage of rDNA methylation as cancer detection markers in plasma.

In present study, we examined methylation pattern of each CpG site in rDNA for tumors and normal tissues with WGBS and RRBS data and found that IGS of rDNA has high methylation level and low variation in normal tissues and plasma samples, but shows hypo-methylation in at least 4 cancer types compared with normal tissues. Finally, we supported that aberrant methylation of CpG sites in rDNA could be used as competitive and robust markers for cancer detection and monitoring cancer progression in plasma.

## Results

### Aberrant methylation of rDNA in malignant tumors

In adult human tissues, the most abundant DNA methylation occurs in cytosine that belongs to cytosine-phosphate-guanine (CpG) dinucleotides and is crucial to various biological processes[43, 44]. Therefore, we only focused on the methylation of CpG sites on rDNA. The reference of human rDNA repeated unit contains 3288 CpG sites in total. In order to coherently describe regions on rDNA, we defined 5 zones according to methylation patterns (see Methods) where Zone 1 occupies rDNA 1-15455 base pairs (bps); Zone 2 occupies 15456-27705bps; Zone 3 occupies 27706- 29080bps; Zone 4 occupies 29081-41755bps; Zone 5 occupies 41756-42999bps (Fig. 1A). Zone 1 is composed of transcribed region and transcription terminator region which is downstream of 3’ external transcribed spacer (ETS). Zone 3 corresponds to the peak of H3K4me1/2/3 and H3K9ac within IGS reported by previous studies[41, 42]. Zone 5 overlaps with rDNA promoter region. Zone 2 and Zone 4 locate within IGS. The distribution of CpG sites in rDNA is not uniform. Analysis of CpG density in rDNA indicated that Zone 1 and Zone 5 have higher CpG density than other regions in rDNA (Fig. 1B).

**Figure 1.**
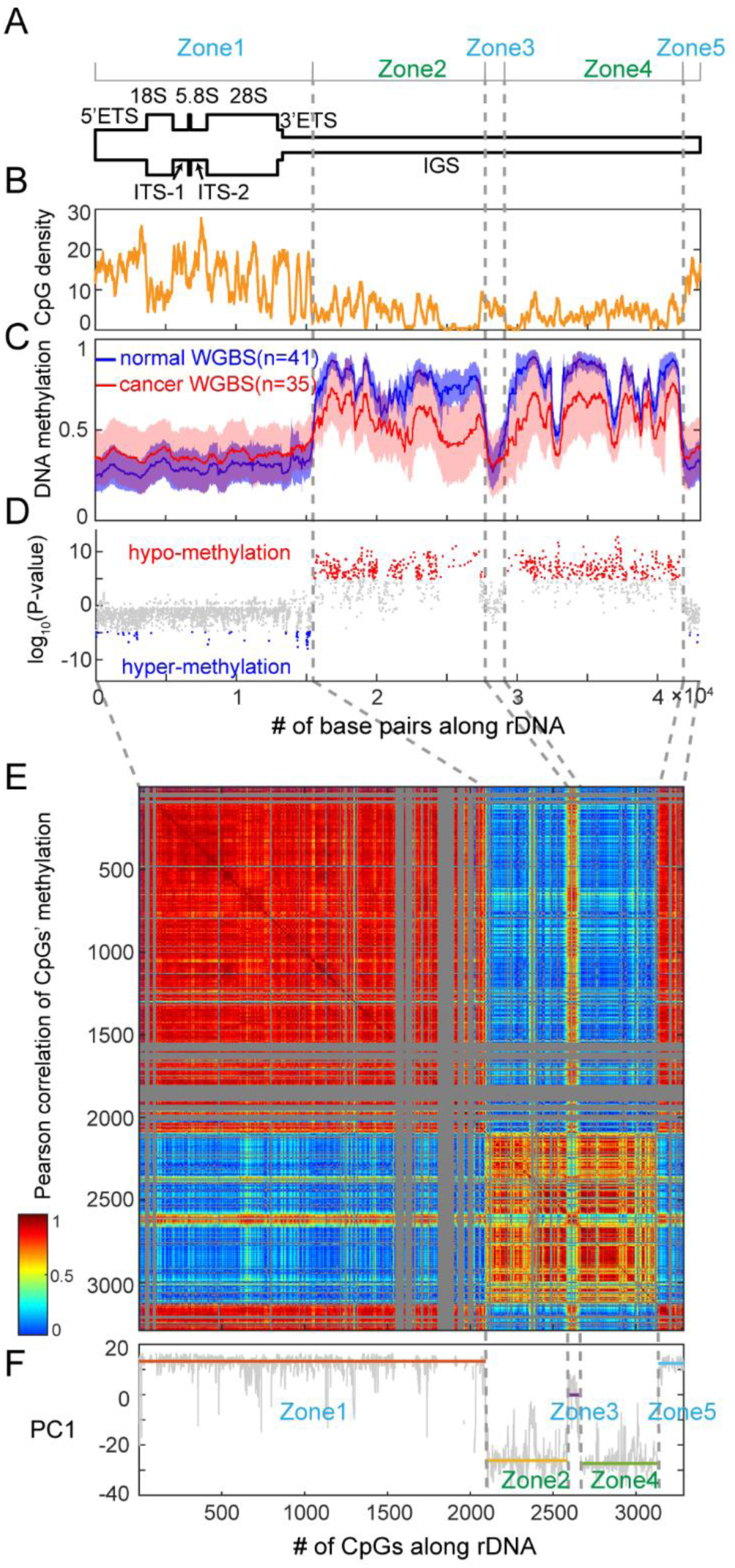
Methylation patterns of tumors and normal tissues. (A) A schematic representation of human rDNA repeat unit. ETS: external transcribed spacer; ITS: internal transcribed spacer; IGS: intergenic spacer. Zones are labeled on the top. (B) Plot of CpG dinucleotides’ density along reference sequences. The vertical axis shows the number of CpG dinucleotides per 100 bps. (C) Plots of methylation levels for normal tissues and tumors. The lines indicate the mean methylation levels of tumors and normal tissues. Scopes of plots show the methylation intervals of ±1 standard derivation. (D) Scatter plot of differential significance between tumors and normal tissues for each CpG along rDNA. Logarithms of P values of Wilcoxson test are shown as ordinate. Positive values mean hypo-methylation in tumors; negative values mean hyper-methylation in tumors. CpG sites of which Bonferroni adjusted P values are smaller than 0.05 are colored as red (hypo-methylation in cancer) and blue (hyper-methylation in cancer). (E) Heat map of correlation matrix between CpG sites along rDNA. Correlation scores of CpG sites with low coverage are labeled as grey. grey dashed lines shows the correspondence between positions and ranked numbers of CpG dinucleotides. (F) Plot of principal component 1 (PC1) of correlation matrix. For each zone, median of PC1 values is drawn as horizontal line.

First, we investigated the general methylation patterns in 45 tumor samples and 41 normal tissue samples using WGBS or RRBS data collected from several studies (see Methods). In normal tissues, Zone 1, 3, 5 have low methylation levels and higher variations, while Zone 2, 4 are relatively highly methylated and less variable (Fig. C, Additional file 1: Fig. S1B). Two trends of methylation pattern changes were observed in malignant tumors. Zone 1, 3, 5 has higher methylation in tumors than normal tissues (Fig. 1C). Zone 2 and 4 within IGS show hypo-methylation in tumors. Further, methylation variations in Zone 2 and 4 increase in tumors (Additional file 1: Fig. S1C). There are more CpG sites that show significantly hypo-methylated than these are significantly hyper-methylated (Fig. 1D).

In addition, Correlation between methylation of CpG sites along rDNA suggests that the CpG sites can be grouped into two categories. CpG sites of Category 1 and Category 2 locate at Zone 1, 3, 5 and Zone 2, 4 respectively. The methylation level of CpG sites has high correlations within the same category but low correlations between different categories. Principal component analysis of correlation matrix shows consistent results of two categories (Fig. 1F).

Above all, through comparison of rDNA’s methylation in normal tissues and tumors, we showed distinct methylation pattern in tumors that might be used as biomarkers of cancer detection.

### Hypo-methylation of IGS is a common feature in different cancer types

In order to further interrogate the features of aberrant methylation in tumors, we divided samples into different subsets and investigated their methylation profiles separately. First, we examined methylation heterogeneity in different normal tissues by using WGBS data set from ten adult tissues of one human donor [9]and found that Zone 2 and 4 show lower methylation variation than Zone 1, 3 and 5 (Additional file 1: Fig. S1D). Further, Methylation profiles of buff coat from 15 individuals show same trends (Additional file 1: Fig. S1E). These results indicate that methylations of rDNA IGS region is more stable across tissue types and individuals, which means differences occurred in IGS can be easier to be detected

Based on the findings in normal tissues, we were curious about whether IGS’s hypo-methylation is a common feature for different cancer types. The majority of IGS (Zone 2 and 4) shows hypo-methylation in at least 4 types of cancer compared with corresponding normal tissues (Fig. 2B-E). In contrast, hyper-methylation in Zone 1, 3 and 5 is not consistent in different cancer types (Fig. 2B-E).

**Figure 2.**
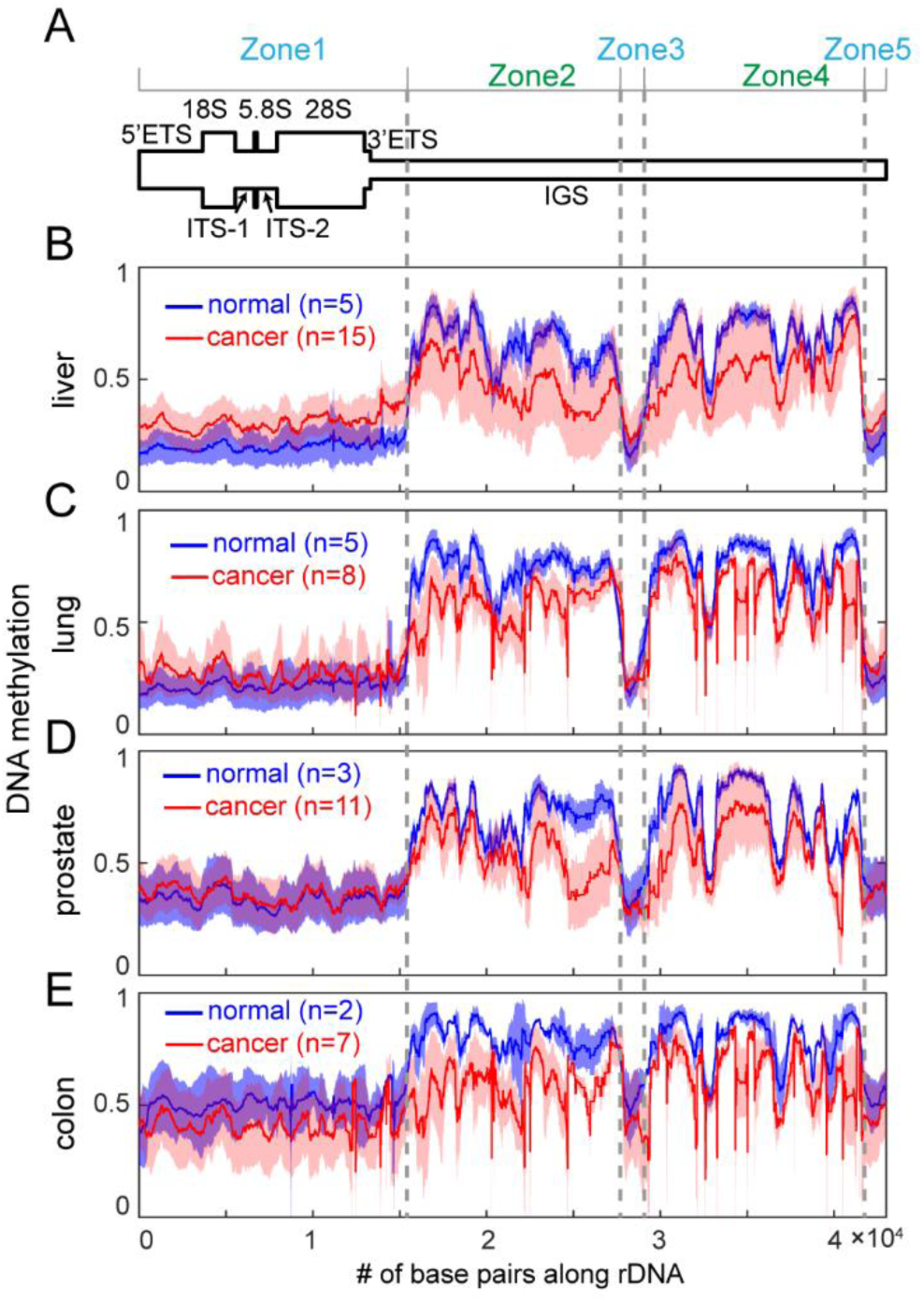
methylation levels in rDNA for 4 cancer types. (A) A schematic representation of human rDNA repeat unit. (B-E) Plots of methylation levels along rDNA reference sequences for four different cancer types and corresponding normal tissues. The red and blue lines are the mean methylation levels of tumors and normal tissues. Scopes of plots show the methylation intervals of ±1 standard derivation.

In summary, Zone 2 and 4 which locate in IGS show low methylation variation in normal tissues and significant differential methylation levels in 4 types of cancer. Because plasma cfDNA is a mix of cfDNA that is released from multiple tissues of human body, low methylation variation of IGS may mean stable signals in healthy plasma, which is benefit for detecting cancer. In following sections, we would step forward to investigate the methylation in plasma cfDNA and examined the possibility of rDNA’s IGS being used as cancer detection markers.

### IGS of rDNA showed hypo-methylation in plasma samples from cancer patients

In liquid biopsy, cfDNA from plasma is one of the most representative analysts which has been widely studied. Plasma cfDNA of healthy subjects comes from various tissues but predominately from blood cells [45, 46]. Robust and accurate detection of cancer in plasma highly depends on the mix ratio of cancer signals and the stability of markers.

At first, Methylation of plasma cfDNA from healthy individuals showed that Zone 1, 3 and 5 have relatively low methylation level and high methylation variation, while Zone 2 and 4 have high methylation level and very low variations, which is similar to normal tissues (Fig. 3B-C). Further, the top ranked differential CpG sites between plasma from healthy subjects and cancer patients are more significant than random shuffling background (Additional file 1: Fig. S2A), which suggests that such differences between healthy subjects and cancer patients are not random noises.

**Figure 3.**
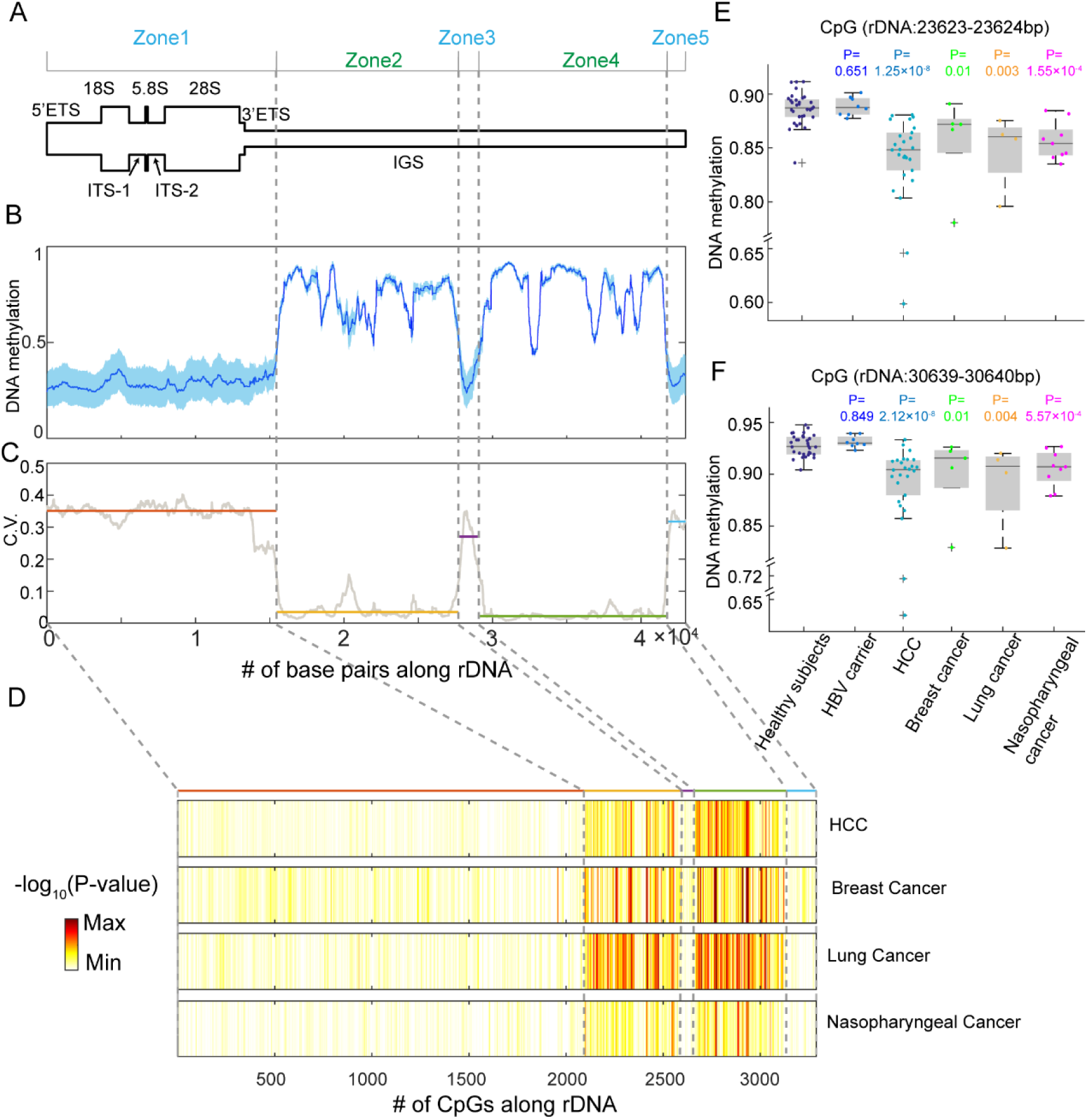
rDNA’s methylation of plasma cfDNA. (A) A schematic representation of human rDNA repeat unit. (B) Plots of methylation levels for plasma cfDNA of healthy subjects. The line indicates the mean methylation level of healthy plasma. Scopes of plots show the methylation intervals of ±1 standard derivation. (C) Plot of coefficient of variance (C.V.) of healthy plasma in rDNA. For each zone, median of C.V. is drew as horizontal line. (D) Heat maps of significance of differences in distinguishing healthy and cancer patients on single CpG dinucleotides. CpG dinucleotides are ranked by positions on rDNA. grey dashed lines show the correspondence between positions and ranked numbers of CpG dinucleotides. (E-F) Boxplots of methylation of two examples of CpG sites at rDNA (23623-23624 bp, 30639-30640) in different plasma samples. P values of Wilcoxon test are labeled on the top of samples.

Analysis also showed that the top significant candidate markers are mainly located in Zone 2 and Zone 4 of IGS in four types of cancer (Fig. 3D). Plasma from cancer patients also showed hypo-methylation in Zone 2 and Zone 4 (Additional file 1: Fig. S3A), which shows the consistent direction of changes with tumor tissues. Two examples of CpG sites which locate at Zone 2 and 4 show hypo-methylation in cancer plasma samples, while there are no significant differences between HBV carriers and healthy subject (Fig. 3E-F).

Further, we interrogated differential CpG dinucleotides throughout comparison between healthy subjects and lung, colon cancer with plasma RRBS data. Though RRBS data prefer genomic regions of high CpG density, plasma from patients of lung cancer and colon cancer also show hypo-methylation at Zone 2 and 4 (Additional file 1: Fig. S3A). And the top ranked significant CpG sites show higher differences than random shuffling (Additional file 1: Fig. S2B). This indicated that hypo-methylation of CpG sites within IGS is a universal pattern in different cancer types. Methylation levels of CpG sites on IGS can serve as markers for cancer detection in plasma.

### rDNA methylation as markers for cancer detection

In order to evaluate the performances of methylation of rDNA in cancer prediction, we randomly divided the datasets into training and testing sets with a ratio of 2:1, and trained classifiers on training dataset and then evaluated the performances on testing dataset. The operation was repeated for 50 times independently to avoid random effects (see Methods). Four classifiers (L1 regularized logistic regression classifier, support vector machine (SVM) classifier, k-nearest neighbor (KNN) classifier and random forest classifier) were used for evaluation.

Since there is only small amount of plasma WGBS samples for lung cancer, breast cancer and nasopharyngeal cancer (see Additional file 2), we performed training and testing only with liver cancer WGBS. For plasma RRBS, we performed the operation on both lung cancer and colon cancer. At first, prediction performance is affected by the number of CpG sites used as markers. The mean of AUCs could achieve 0.85 with only one CpG site, and 0.92 with two CpG sites using liver plasma WGBS (Fig. 4A). The prediction performance becomes stable when the number of CpG sites is 10. Similar result was observed on plasma RRBS (Additional file 1: Fig. S4A). Therefore, we chose 10 CpG sites as markers of classifiers in following study. The CpG sites selected by machine learning mainly located in Zone 2 and 4 of IGS (Additional file 3-5).

**Figure 4.**
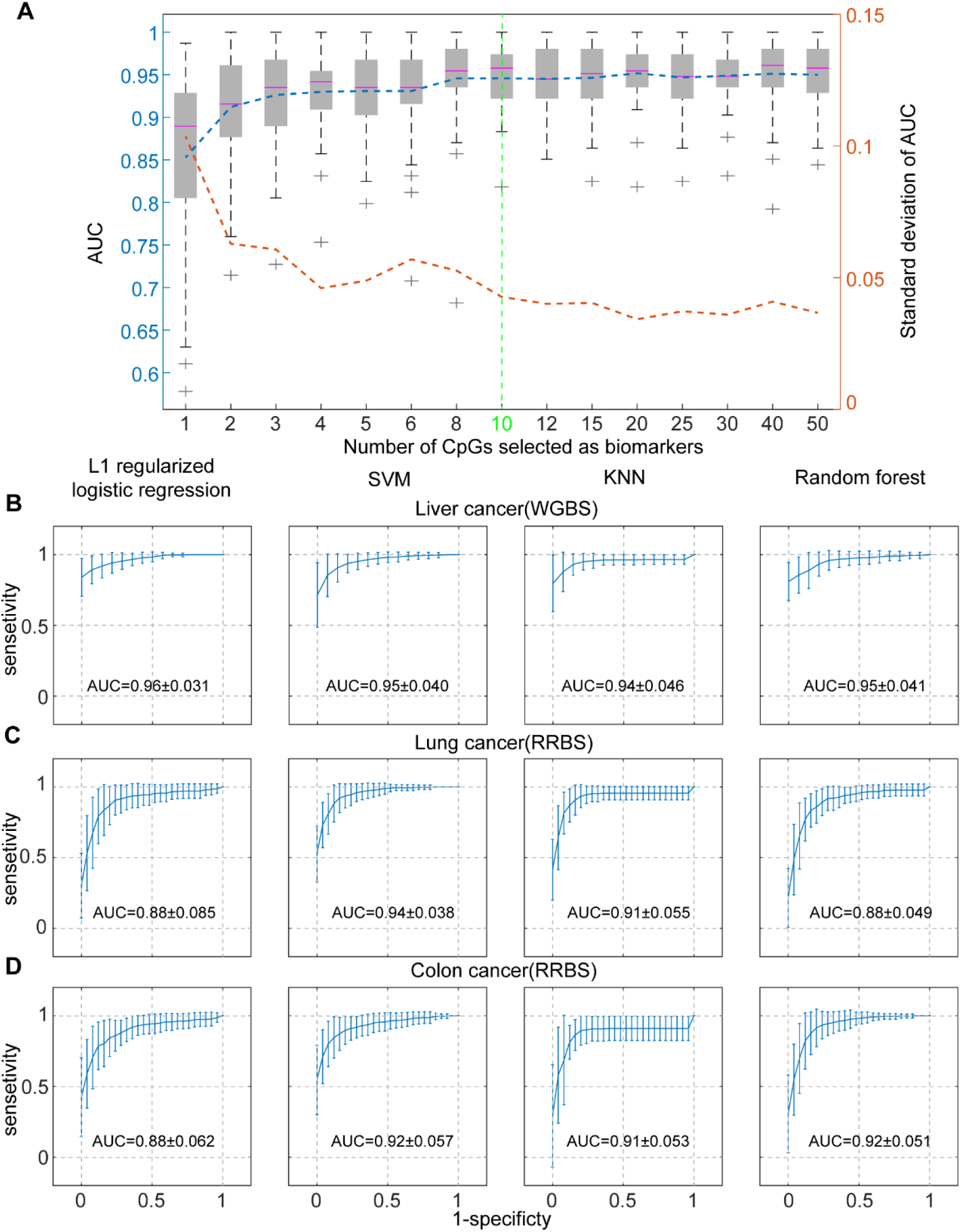
Performances in distinguishing healthy and liver cancer plasma. (A) Boxplots of AUCs of SVM classifier with different numbers of CpG sites. Blue dashed line shows mean AUCs, while red dashed line shows AUCs’ standard derivation. (B) Receiver operating characteristic (ROC) curves of four classifiers for liver plasma WGBS testing set. Standard derivation bars are labeled with the performance variations of 50 runs. Names of classifiers are labeled on the top of ROC curves. AUCs are labeled on the plots with format of mean±1 standard derivation. (C) ROC curves of four classifiers for lung plasma RRBS testing set. (D) ROC curves of four classifiers for colon plasma RRBS testing set.

Training area under curves (AUCs) are almost 1 and testing AUCs are more than 0.94 in liver cancer with all four classifiers (Additional file 1: Fig. S5A, Fig. 4B). rDNA methylation shows competitive performance in liver cancer detection compared with alpha-fetoprotein (AFP) which was widely used in clinical with AUC of 0.80[47], six-DNA methylation-marker panel with AUC of 0.94[47] and whole genome hypo-methylation method with AUC of 0.93[8]. Similar to liver cancer WGBS, four models training on lung cancer RRBS and colon cancer RRBS showed fairly good performances both in training and testing sets compared with original report of this dataset [9] (Additional file 1. S5B-C, Fig. 4C-D). For example, testing AUCs achieved 0.94 and 0.92 respectively (Fig. 4C-D). Above all, these results suggested that methylation of IGS in rDNA could predict whether a patient has cancer with high accuracy.

Although methylation of rDNA showed high prediction performance in single cancer type, we wondered whether CpG sites for cancer detection could transfer from one cancer type to others. Classifiers trained with one cancer type predicted effectively for other cancer types within WGBS or RRBS data set (Additional file 1: Fig. S6A-C). Especially in RRBS dataset of lung cancer and colon cancer, models trained with the other cancer type could obtained similar performances as models trained with the same cancer type data (Additional file 1: Fig. S6B-C). And the biomarkers selected using WGBS showed good prediction performance on RRBS data, and vice versa (Additional file 1. S7A-D, 8A-D). However, when we trained models to distinguish lung cancer samples from colon cancer samples with RRBS data, the performance was poor (Additional file 1: Fig. S9A-B). These results suggested that methylation of plasma rDNA might act as potential markers for pan-cancer detection.

### Robustness of rDNA methylation as cancer detection markers

We have mentioned that one great advantage of rDNA for acting as cancer-detection markers is its high copy number. High copy number may enhance the robustness of signal. WGBS covers whole genome and is expensive, so we examined whether methylation of rDNA could keep its prediction performance after down-sampling WGBS sequencing reads. Plasma WGBS of liver cancer was sequenced for one lane (mean 163 million (M) raw reads) of an Illumina HiSeq2000 sequencer per sample, we down-sampled raw sequencing reads to 100M, 50M, 20M and 10M, then trained and tested with down-sampled raw sequencing reads (see Methods).

Four classifiers showed slightly decreased performances with down-sampled reads (Fig. 5A). However, even with 10 million raw reads (0.25× whole genome coverage), four classifiers could obtain a mean AUC of at least 0.90. L1 regularized logistic regression and random forest classifiers’ mean AUC could reach 0.93. It indicated that rDNA methylation acting as cancer detection markers is robust. Robustness of rDNA methylation means that we can reduce the sequencing depth of WGBS to reduce cost of sequencing for following studies or clinical applications.

**Figure 5.**
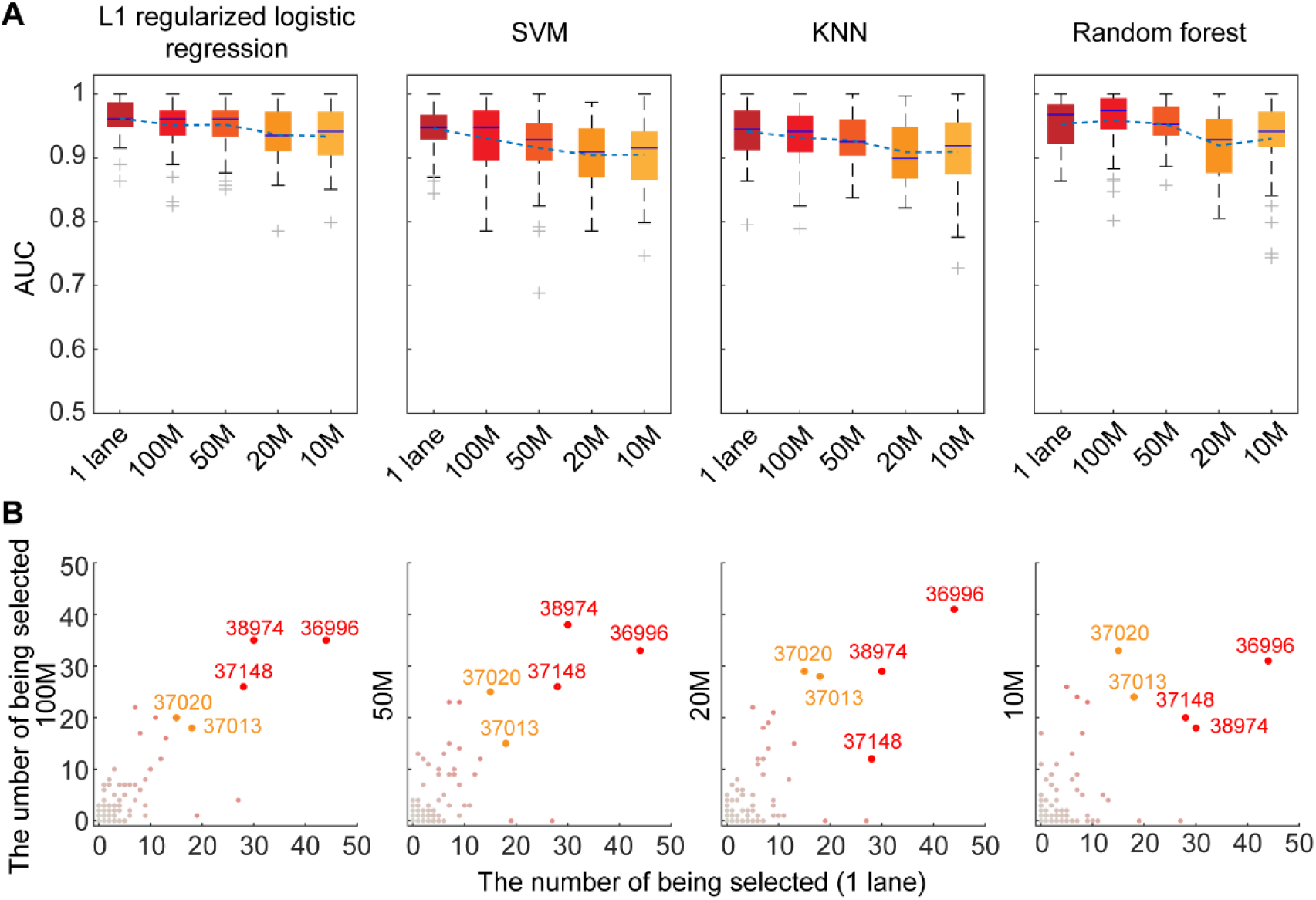
Prediction performances based on down sampling sequencing reads. (A) Boxplots of AUCs in different down sampling depths for four models. Horizontal axis shows the number of raw reads after down sampling. 1 lane is the original of sequencing depth (mean depth: 163 million). Models are trained and tested with corresponding down-sequencing depths. (B) Scatter plots of the number of being selected as markers for each CpG site in different sequencing depths. Horizontal axis of four scatter plots is the number of being selected for each CpG site with 1 lane sequencing depth. Ordinate is the number of being selected for each CpG site with 100M, 50M, 20M and 10M sequencing depths. Red dots indicate three CpG sites with high selected numbers in both high and low sequencing depths. Orange dots shows two CpG sites with higher selected number in low sequencing depth. The first position of each CpG site in rDNA is labeled around CpG dot.

The number of being selected as markers were counted for each CpG site in feature selection. With different sequencing depths, CpG sites which were frequently selected as markers were consistent (Fig. 5B). CpG sites at rDNA: 36996-36997bps, rDNA: 38974-38975bps, rDNA: 37148-37149bps showed high frequencies of being selected in both high and low sequencing depths. CpG sites at rDNA: 37020-37021bps and rDNA: 37013-37014bps showed much higher frequencies in low sequencing depth. CpG sites being selected in down-sampling dataset mainly locate at Zone 2 and 4 of IGS.

In summary, methylation of IGS in rDNA serves as robust markers for cancer detection in plasma.

### Methylation of rDNA acting as markers for monitoring cancer progression and treatment

We also investigated whether methylation of rDNA could be used for monitoring the cancer progression and surgery. There were plasma WGBS of time series from two liver cancer patients, which included plasma WGBS before and after tumor resection. L1-regularized logistic regression classifier trained with liver cancer plasma samples was used to evaluate the effects of surgery. The patient TBR34 died of metastatic disease 8 months after surgery[8], which was consistent with the prediction that TBR34 still had high probability of having cancer after surgery (Fig. 6A). The patient TBR36 was still alive beyond 20 months after surgery[8], which was consistent with the fact that the predicted probability of having caner decreased obviously 3 days after surgery (Fig. 6A).

**Figure 6.**
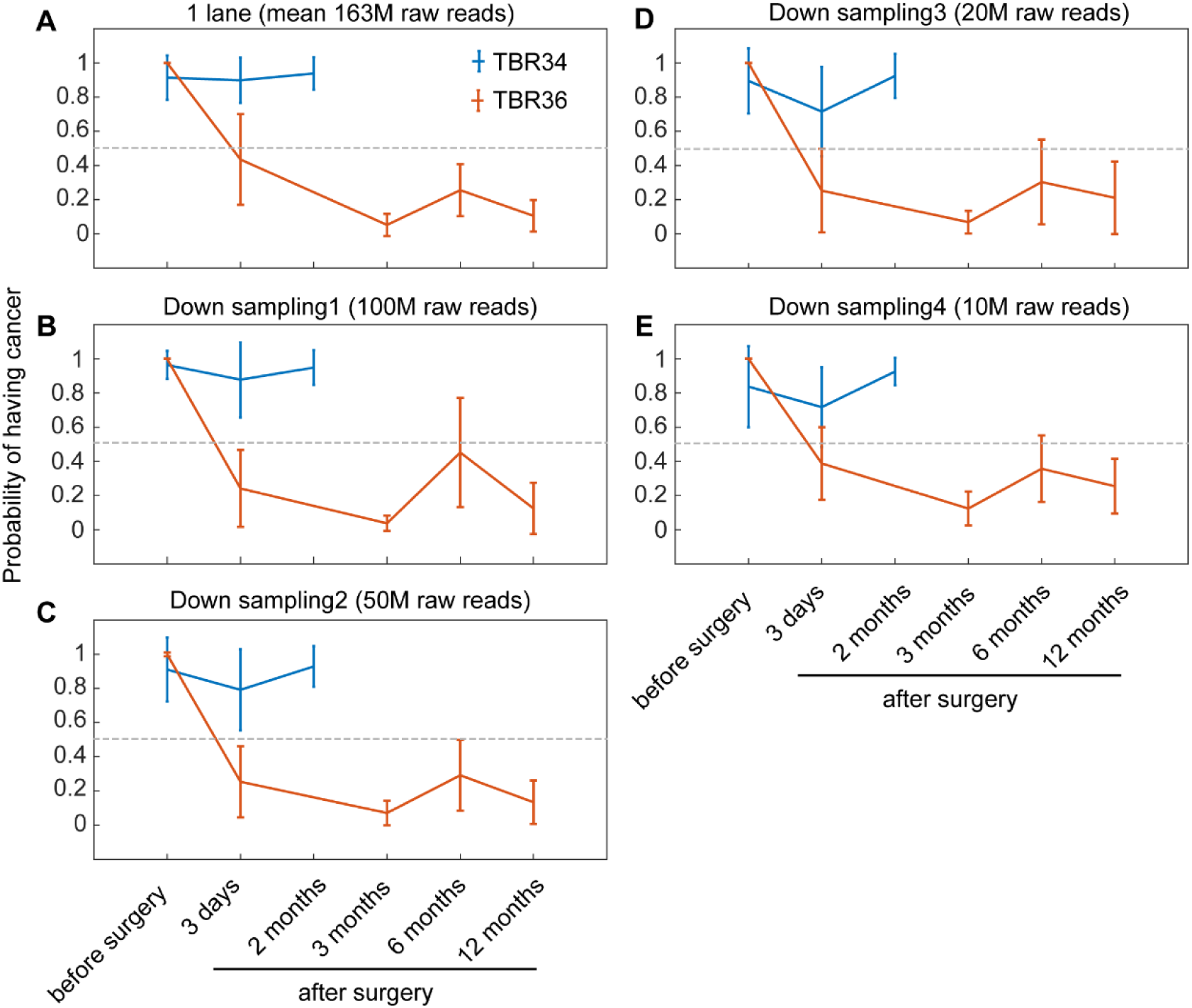
Monitoring cancer progression for two patients. (A) Bar boxplots of predicted probabilities of having cancer for patient TBR34 and TBR36. Horizontal axis shows the time lines of tumor resection. Horizontal grey line shows the threshold of 0.5. (B-E) Bar boxplots of predicted probabilities of having cancer for two patients based on different down sampling depths. The predicted models are trained and tested in corresponding down-sequencing depths.

Further, performance of down-sampling sequence reads was checked and the prediction results in different down-sampling sequencing depths were similar to that of no down-sampling data (Fig. 6B-E). The results mean that we can evaluate the surgery effects with low sequencing depth and take measures ahead of cancer progression. Taken together, methylation of rDNA can also be used as robust markers for monitoring cancer progression and treatment.

## Discussion

In this study, we used WGBS and RRBS data to investigate the methylation patterns of rDNA at each single CpG site for multiple cancer types and identified the aberrant methylation patterns of cancer. We found that CpG sites within IGS are promising markers for cancer detection. These CpG sites have high methylation level and low variation both in normal tissues and plasma, while have decreased methylation levels in four cancer types. We evaluated the performance of these markers with plasma bisulfite sequencing samples from healthy subjects and cancer patients. Further, we found these markers could also be used for monitoring cancer progression and treatment.

In this study, we showed that hypo-methylation of reduced IGS is a universal feature in 4 cancer types, which was rarely reported before. In previous studies, researchers kept focusing on methylation changes of promoter or transcription body of rDNA. In breast cancer, ovarian cancer and endometrial carcinoma, hyper-methylation of transcribed domain of rDNA were reported as an important feature and its methylation level was correlated with patients’ survival rates[37–39]. However, this feature can hardly be used as markers in plasma biopsy because methylation of these regions has high variation in both plasma samples of healthy subjects and normal tissues. In contrast, CpG sites in IGS will be attractive biomarkers for both solid biopsy and liquid biopsy.

Using rDNA methylation as markers for cancer detection has one obvious advantage – robustness. Studying the methylation pattern of CpG sites in non-repeated genome regions required very high coverage to get stable signals. In this study, down-sampling experiments showed that sequencing depth of 10 million raw reads (0.25× whole genome coverage) could also obtain competitive prediction performance due to high copy number, which provides us a robust biomarker. In addition to non-targeted bisulfite sequencing, designing probes or primers that target the differential CpG sites in rDNA will be an alternative way.

However, there were some limitations in this study, which need to be optimized for better outputs with the developments of technologies and databases. First, sequencing reads were mapped to only one reference rDNA sequence and signals were aggregated for all copies of rDNA in this study. However, it was reported that rDNA sequence has a certain rate of mutations, especially in IGS[13]. Recently, two studies has explored the sequence variation in rDNA[48, 49]. Researchers studied chromosome 21 of one individual and identified 101 and 235 variant positions in transcribed regions and IGS respectively[49]. By analyzing whole genome sequencing data from 1000 Genome Project, researchers also identified intra- and inter-individual nucleotide variations in transcribed regions and found variant ribosomal RNA alleles might be correlated with physiology and diseases[48]. With the development of human genetic variants about rDNA, sequencing reads can be mapped to multiple rDNA alleles, which may assist us to investigate the methylation patterns of rDNA and detect cancer more accurately.

In addition, fragments with low methylation levels are more easy to be damaged during bisulfite treatment in library construction[50]. It means that hypo-methylation of cancer tissues may be more hard to be detected due to bisulfite conversion. If there are alternative technologies which retain the hypo-methylation fragments, it may improve the power of rDNA’s methylation in cancer detection. For example, third generation sequencing can detect DNA methylation without any processing on DNA samples[51].

Finally, methylation of rDNA is a potential marker for pan-cancer detection and can be used for screening potential cancer patients. However, it may need to combine with other cancer type-specific markers to achieve cancer type-specific predictions. It has been suggested that combining multiple parameters or analysts enhances the performance of liquid biopsy[1, 2]. Therefore, methylation of rDNA can serve as perfect partner markers for cancer managements.

## Conclusion

In this study, we provided high-resolution map of DNA methylation at each single CpG site for cancer with bisulfite sequencing data. We found that CpG sites in IGS of rDNA has high methylation level and low variation in both normal tissues and plasma and showed hypo-methylation in 4 cancer types. The study supported that methylation of IGS in rDNA provides a competitive and robust marker for detecting cancer and monitoring cancer progression and treatment.

## Methods

### Data collection

We collected WGBS or RRBS data of tumors, normal tissues and plasma cfDNA from published dataset[8, 9, 52, 53]. In total, we obtained 15 types of normal tissues and 4 types of cancer (see Additional file 2). Cancer types were selected based on that the number of samples from one cancer type is more than 5. In this study, we analyzed 41 samples of normal tissues and 45 samples of tumors and investigated the methylation patterns in rDNA. We also collected WGBS and RRBS data of plasma cfDNA from healthy subjects and patients with 5 different cancer types (see Additional file 2). There are totally 224 plasma cfDNA samples. These data were downloaded from Gene Expression Omnibus (GEO) under accession number GSE52272, GSE70091, GSE79279, GSE104789 and European Genome–Phenome Archive (EGA) under accession number EGAS00001000566.

### Analyses of WGBS and RRBS data

Downloaded data were mapped to human ribosomal DNA repeat unit reference sequences (Genebank: U13369.1, https://www.ncbi.nlm.nih.gov/nuccore/U13369.1) with software BS- Seeker2[54]. We used ‘local alignment’ mode to map these data. Mapped bam files were used to call methylations at single CpG dinucleotides without removing duplications because there are extremely high coverages in rDNA. We only considered the methylation patterns of CpG dinucleotides as there were low methylation levels at non-CpG sites in human genome. We removed mapped reads which had alignment lengths less than 40 bps. Too small alignment length caused higher false positive hitting. Then, the numbers of methylation and un-methylation reads were counted at each single CpG sites. Finally, the methylation levels of each CpG sites were calculated unless the mapped reads number in each sample was less 20.

### Dividing rDNA into different zones

Different regions in rDNA showed different methylation patterns. In order to describe the methylation patterns of rDNA, we divided rDNA into 5 different non-overlapped zones according to methylation of CpG sites. First, we calculated the methylation levels and correlation matrix of CpG sites in normal tissues and tumors. Next, we identified boundaries of CpG sites of different zones roughly. We also calculated PC1 of correlation matrix by performing PCA on correlation matrix. We identified the exact CpG site boundaries by shifting rough boundaries several CpG sites to achieve the largest differences of median PC1 between adjacent zones. Finally, boundary coordinates were determined according to CpG boundaries.

### Training classifiers and predicting

For liver cancer, we randomly divided healthy subjects, HBV carrier and liver cancer patients into training and testing sets with a ratio of 2:1. In training set, we screened the markers and trained classifiers; in testing set, we validated the chosen markers and models trained with training set.

In training set, we drew ROC curve and calculate the AUC for each CpG site in distinguishing cancer patients and non-cancer subjects. The higher the AUC was, the better distinguishing effect was. We sorted the CpG sites according to the AUC scores and chose the top 200 ranked markers. Then we used SVM classifier to select a certain number of features (CpG sites) from top 200 ranked markers with training dataset. In order to make sure that selected differential markers in plasma were due to the differences of liver tumor tissues and normal tissues, we checked the methylation pattern of markers in liver tumor tissues and buff coat. We tried several classification methods to combine the selected markers for predicting whether a person is a cancer patient. These classification methods included L1-regularied logistic regression, supported vector machine, k-nearest neighbor and random forest.

In testing set, we validated the screened markers and trained models by calculating AUCs. In order to avoid random noises, we randomly divided the datasets for 50 times independently and computed the mean AUCs in 50 independent training-testing cases.

For lung cancer, breast cancer and nasopharyngeal cancer which had small number of WGBS samples, we didn’t perform the training and testing operations. For lung cancer and colon cancer which had enough numbers of RRBS samples, the training and testing steps were the similar as liver cancer.

### Down-sampling sequencing reads

In order to test the robustness of markers in rDNA, we down-sampled WGBS data of healthy, HBV carrier and liver cancer plasma samples into different sequencing depths. Each sample of plasma WGBS used one lane of an Illumina Hiseq 2000 sequencer and obtained an average of 163 million raw reads. We down-sampled raw sequencing reads into 100 million, 50 million, 20 million and 10 million subsets. Based on the down-sampled raw reads, we performed the same training and testing steps to study the robustness of rDNA markers. As RRBS data had been enriched signals coming from CpG-rich regions through adjusted biological technology, down-sampling was not applied to RRBS data.

## Supporting information

Additional file 2 Data collection information

Additional file 3 CpG sites selected for liver cancer detection (WGBS)

Additional file 4 CpG sites selected for lung cancer detection (RRBS)

Additional file 5 CpG sites selected for colon cancer detection (RRBS)

## Additional files

Additional file 1: Supplementary Materials

Additional file 2: Data collection information

Additional file 3: CpG sites selected for liver cancer detection (WGBS)

Additional file 4: CpG sites selected for lung cancer detection (RRBS)

Additional file 5: CpG sites selected or colon cancer detection (RRBS)

## Abbreviations

AFP: alpha-fetoprotein
AUC: area under curves
bp: base pair
cfDNA: cell free DNA
CpG: cytosine-phosphate-guanine
ctDNA: circulating tumor DNA
C.V.: coefficient of variance
ETS: external transcribed spacer
IGS: intergenic spacer
ITS: internal transcribed spacer
KNN: k-nearest neighbor
M: million
rDNA: ribosomal DNA
ROC: Receiver operating characteristic
RRBS: reduced representation bisulfite sequencing
SVM: support vector machine (SVM) classifier
WGBS: Whole genome bisulfite sequencing

## Acknowledge

The authors greatly acknowledge Dr. Yuk Ming Dennis Lo and his research group in the Chinese University of Hong Kong for providing the cfDNA WGBS data [8].

## Availability of data and materials

The original datasets analyzed in the current study were reported previously[8, 9, 52, 53] and can be downloaded from Gene Expression Omnibus (GEO) under accession number GSE52272, GSE70091, GSE79279, GSE104789 and European Genome–Phenome Archive (EGA) under accession number EGAS00001000566. The list of accessions and corresponding cancer types in the current study were provided in Additional file 2. The CpG sites which were selected as biomarkers were provided in Additional file 3-5.

## Authors’ contributions

XZ and XW conceived the study and designed the methodological framework. XZ collected, processed the WGBS and RRBS data and performed followed analyses. XZ wrote the manuscript with input from XW, HF, WZ, BZ and YL. All authors read and approved the final manuscript.

## Ethics approval and consent to participate

Not applicable

## Consent for publication

Not applicable

---

**Supplementary Figure 1.**
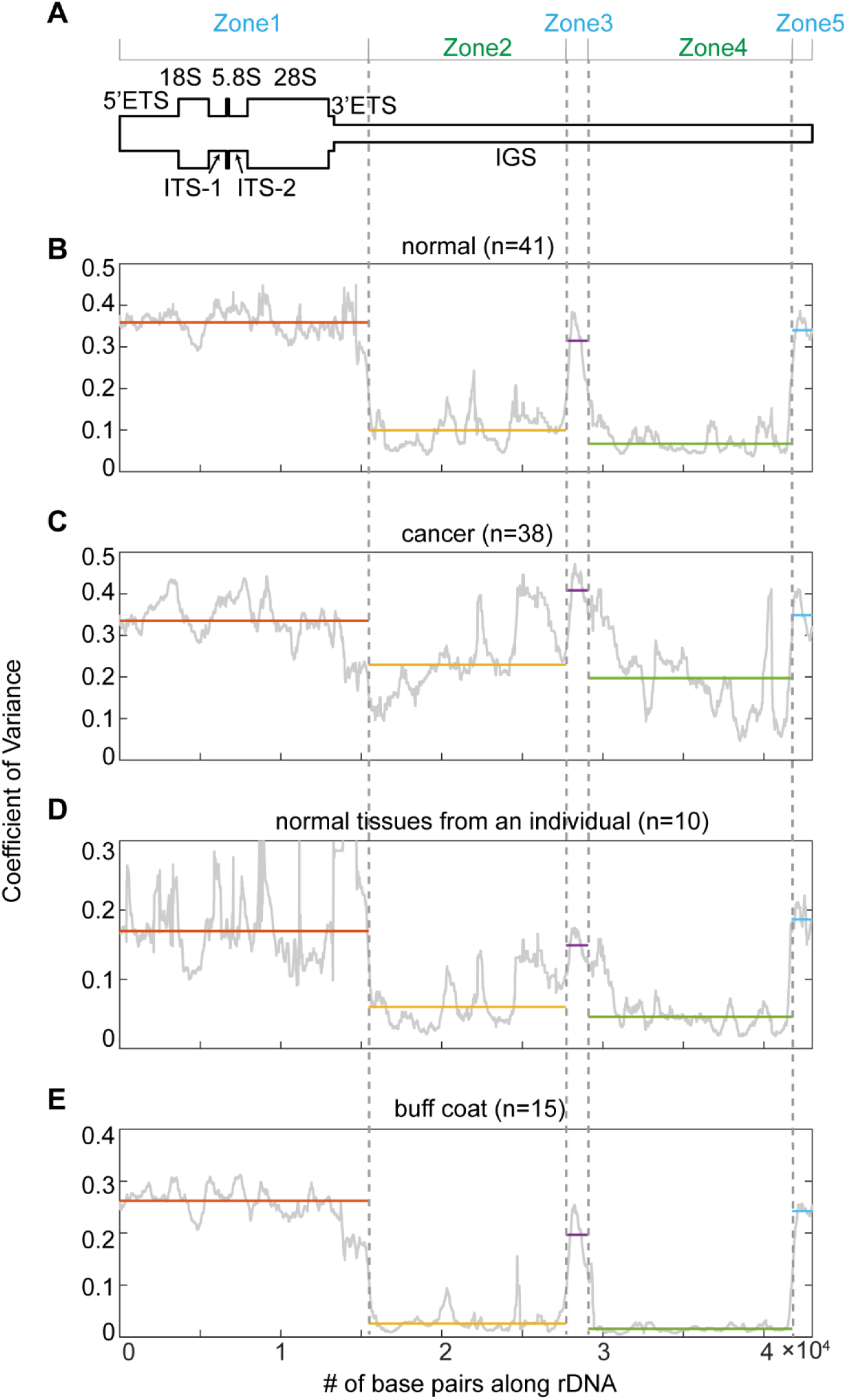
Methylation variations across tumors, normal tissues and individuals. (A) A schematic representation of human rDNA repeat unit. (B) Plot of coefficient of variance (C.V.) of normal tissues in rDNA. For each zone, median of C.V. is drew as horizontal line. (C) Plot of C.V. of tumors on rDNA. (D) Plot of C.V. of ten adult tissues from the same individual on rDNA. (E) Plot of C.V. of buff coats from 15 individuals.

**Supplementary Figure 2.**
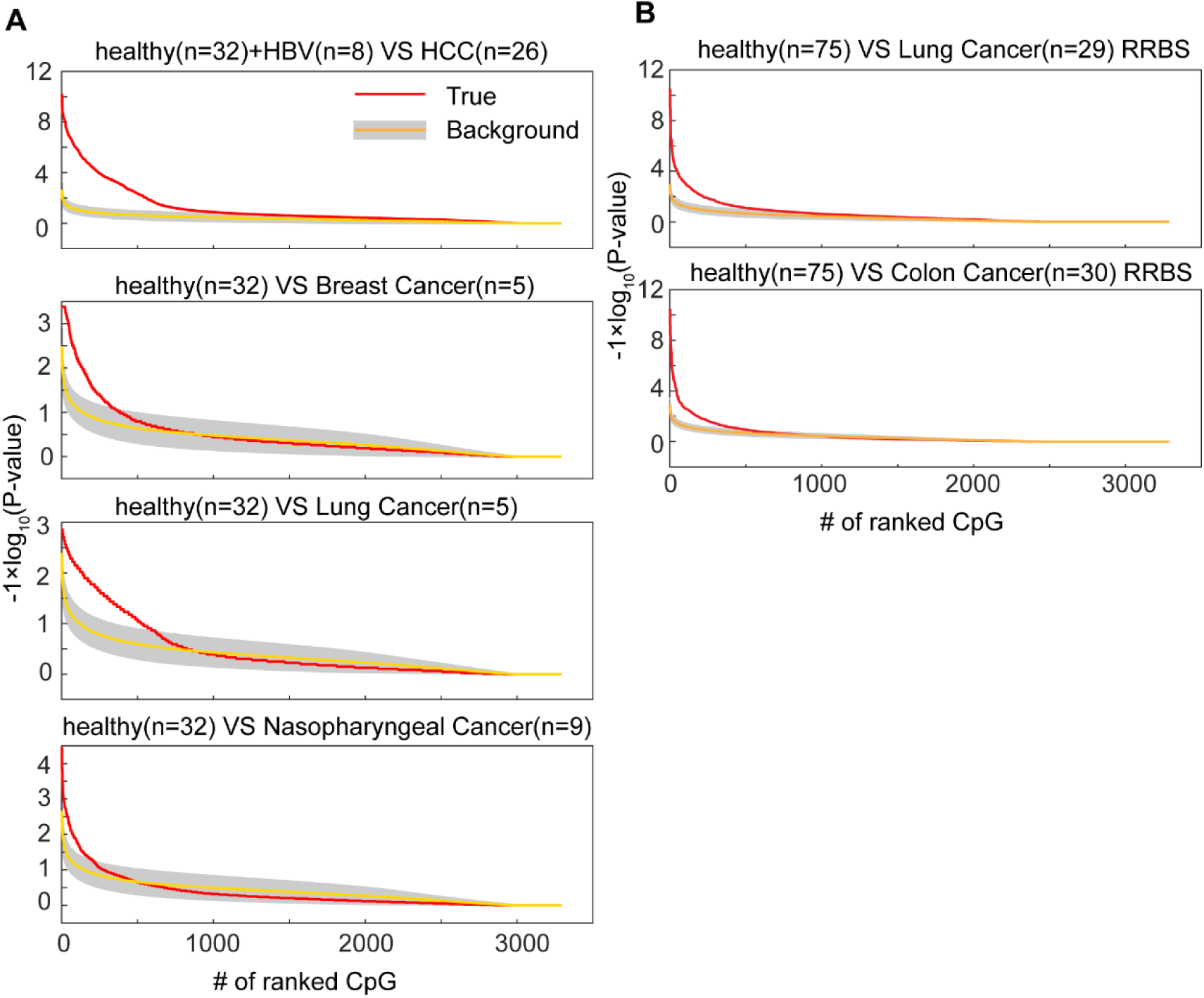
Significance of differences between healthy and cancer plasma. (A) Plots of significance of differences between healthy subjects and cancer patients with WGBS data. CpG dinucleotides are ranked by the P-values of Wilcoxon rank sum test. Red solid lines show the real results; yellow solid lines with grey covers show the results of 100 times’ shuffling. Grey covers shows the intervals of ±1 standard derivation. (B) Plots of significance of differences between healthy subjects and cancer patients with RRBS data.

**Supplementary Figure 3.**
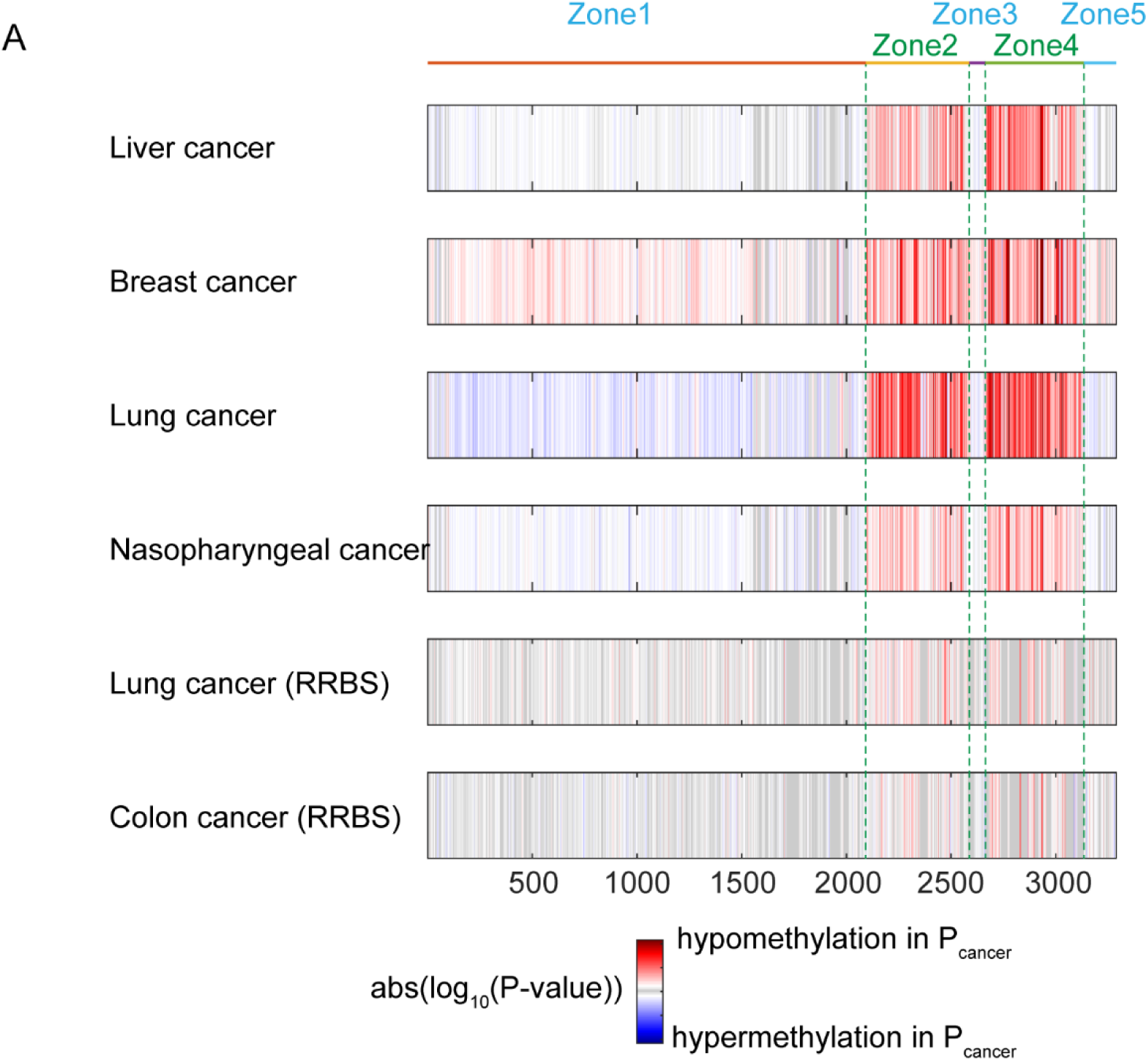
Differential CpG sites between healthy plasma and cancer plasma. (A) Heat maps of significance of differences distinguishing healthy and cancer patients on single CpG dinucleotides. CpG dinucleotides are ranked by positions on rDNA. The maximum of red or blue color indicates the most significant CpG’s absolute log10(P-value) no matter it is either hypo-methylation or hyper-methylation. On the heat maps, we found that almost all of significant CpG sites show hypo-methylation.

**Supplementary Figure 4.**
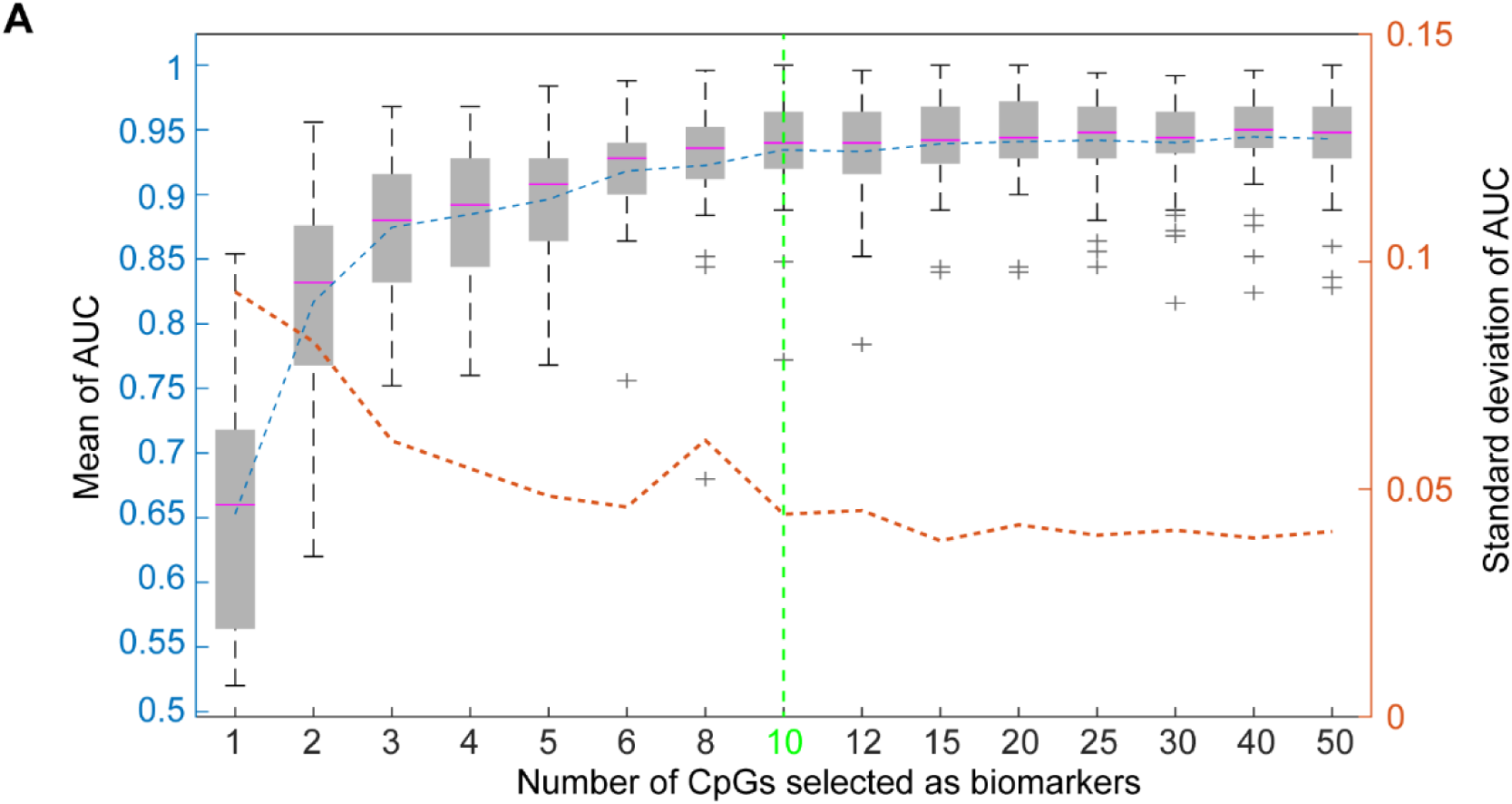
Performances in distinguishing healthy and lung cancer plasma with different numbers of CpG sites (RRBS) (A) Boxplots of AUCs of SVM classifier with different numbers of CpG sites. Blue dashed line shows mean AUCs, while red dashed line shows AUCs’ standard derivation.

**Supplementary Figure 5.**
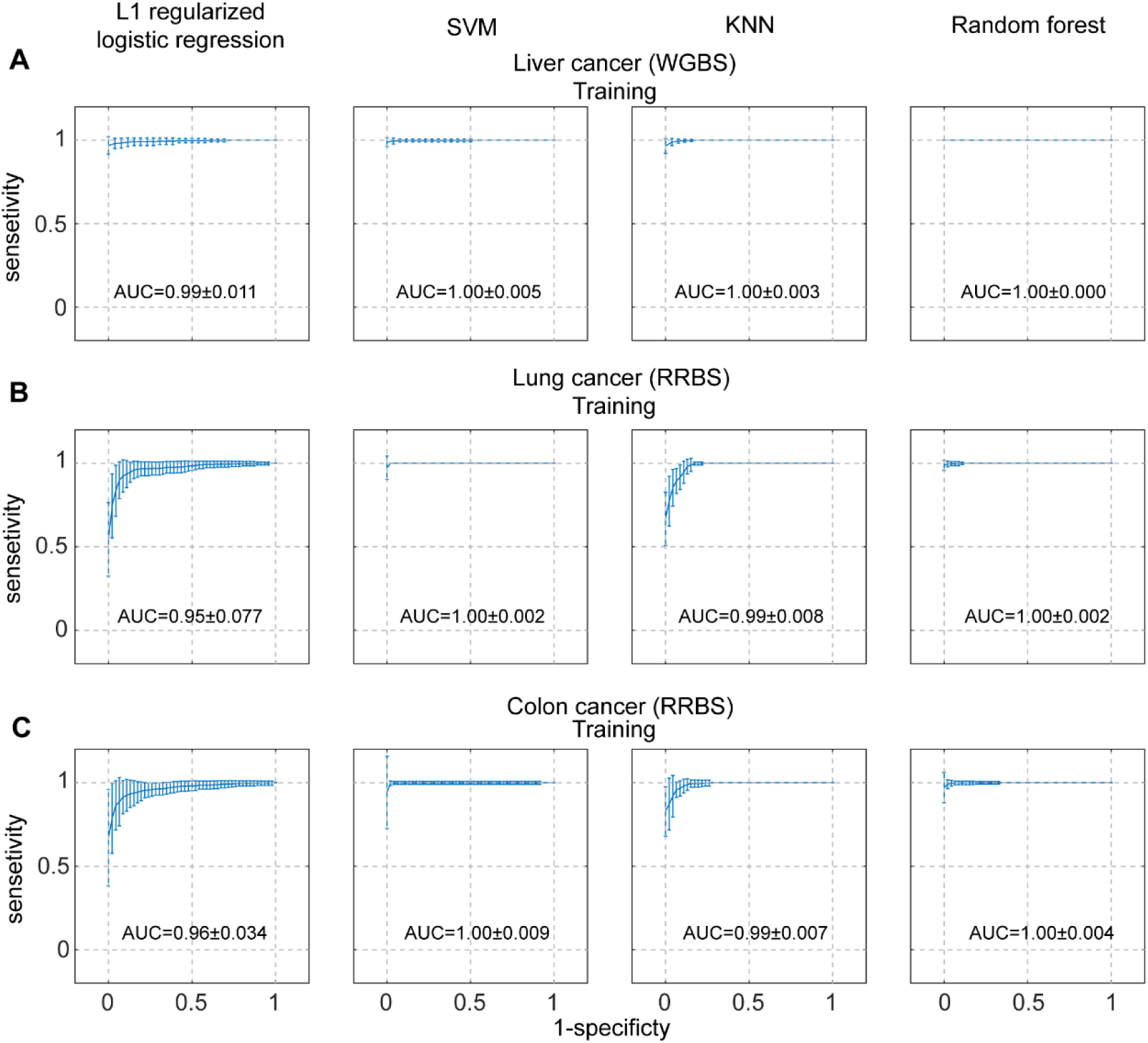
Prediction performances of training dataset. (A) Training ROC curves for liver cancer plasma and healthy plasma with four classifiers (WGBS). Standard derivation bars are labeled showing the performance variation of 50 runs. AUCs are labeled on the plots. (B) Training ROC curves for lung cancer plasma and healthy plasma with four models (RRBS). (C) Training ROC curves for colon cancer plasma and healthy plasma with four models (RRBS).

**Supplementary Figure 6.**
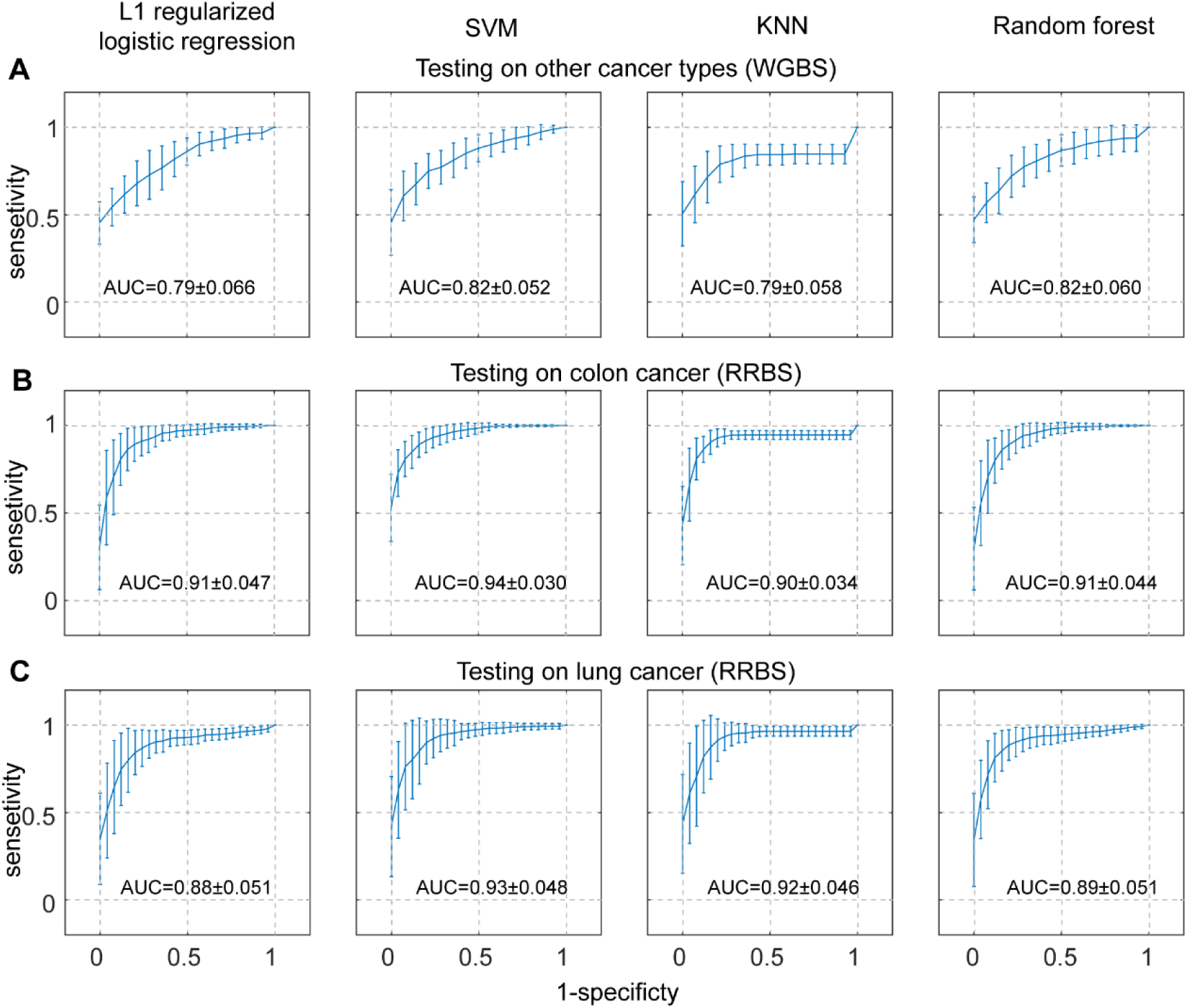
Prediction performances of other cancer types. (A) Testing ROC curves for other cancer types (including breast cancer, lung cancer and nasopharyngeal cancer) plasma WGBS data with four classifiers trained with liver cancer plasma WGBS. Standard derivation bars are labeled showing the performance variation of 50 runs. AUCs are labeled on the plots. (B) Testing ROC curves for colon cancer plasma RRBS with four classifiers trained with lung cancer plasma RRBS. (C) Testing ROC curves for lung cancer plasma RRBS with four classifiers trained with colon cancer plasma RRBS.

**Supplementary Figure 7.**
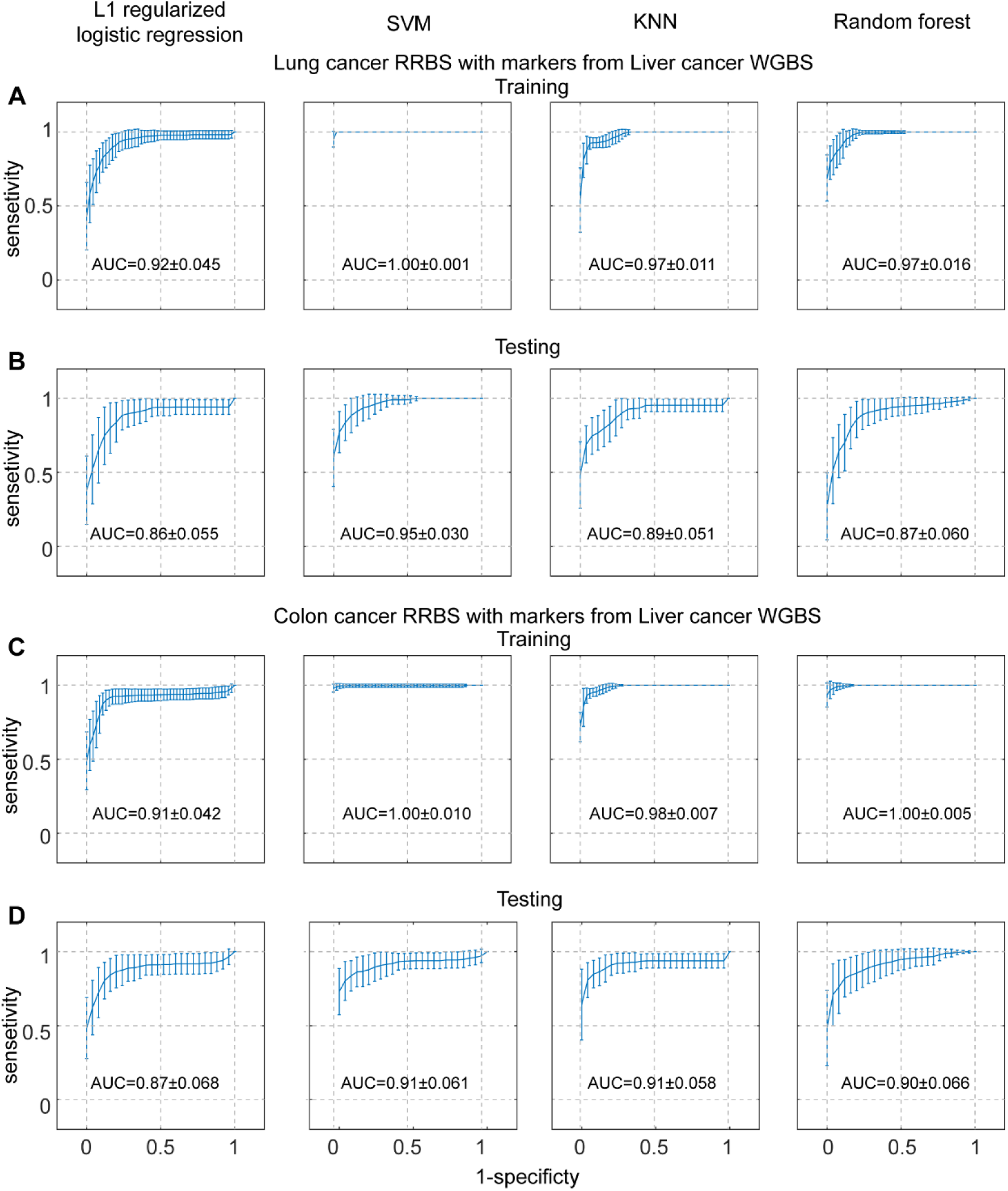
Prediction performances of RRBS plasma with CpG sites transferred from liver cancer WGBS dataset. (A) Training ROC curves for lung cancer plasma RRBS with CpG sites selected in liver cancer WGBS dataset. Standard derivation bars are labeled showing the performance variation of 50 runs. AUCs are labeled on the plots. (B) Testing ROC curves for lung cancer plasma RRBS with CpG sites selected in liver cancer WGBS dataset. (C) Training ROC curves for colon cancer plasma RRBS with CpG sites selected in liver cancer WGBS dataset. (D) Testing ROC curves for colon cancer plasma RRBS with CpG sites selected in liver cancer WGBS dataset.

**Supplementary Figure 8.**
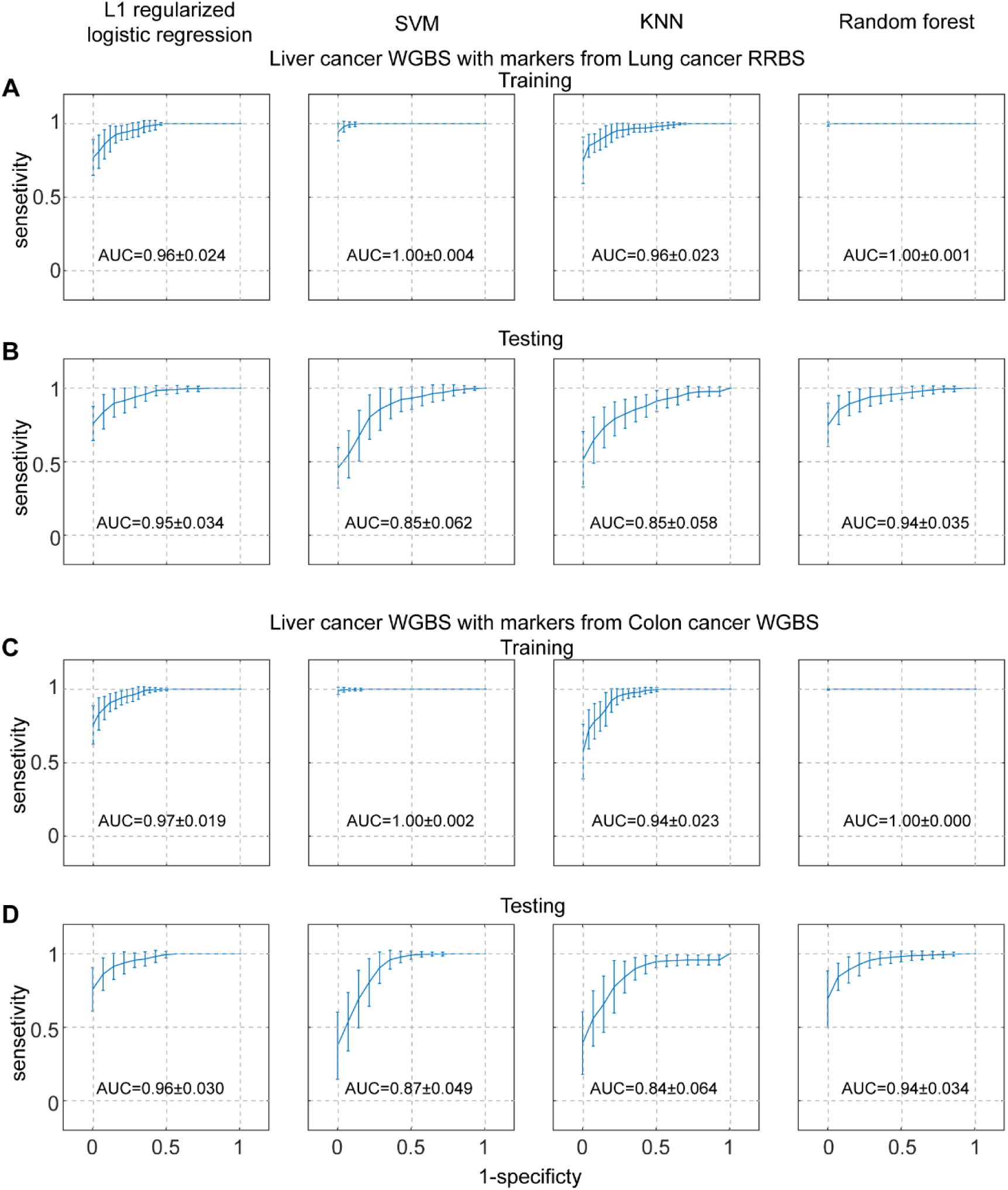
Prediction performances of liver cancer plasma WGBS with CpG sites transferred from RRBS dataset. (A) Training ROC curves for liver cancer plasma WGBS with CpG sites selected in lung cancer plasma RRBS dataset. Standard derivation bars are labeled showing the performance variations of 50 runs. AUCs are labeled on the plots. (B) Testing ROC curves for liver cancer plasma WGBS with CpG sites selected in lung cancer plasma RRBS dataset. (C) Training ROC curves for liver cancer plasma WGBS with CpG sites selected in colon cancer plasma RRBS dataset. (D) Testing ROC curves for liver cancer plasma WGBS with CpG sites selected in colon cancer plasma RRBS dataset.

**Supplementary Figure 9.**
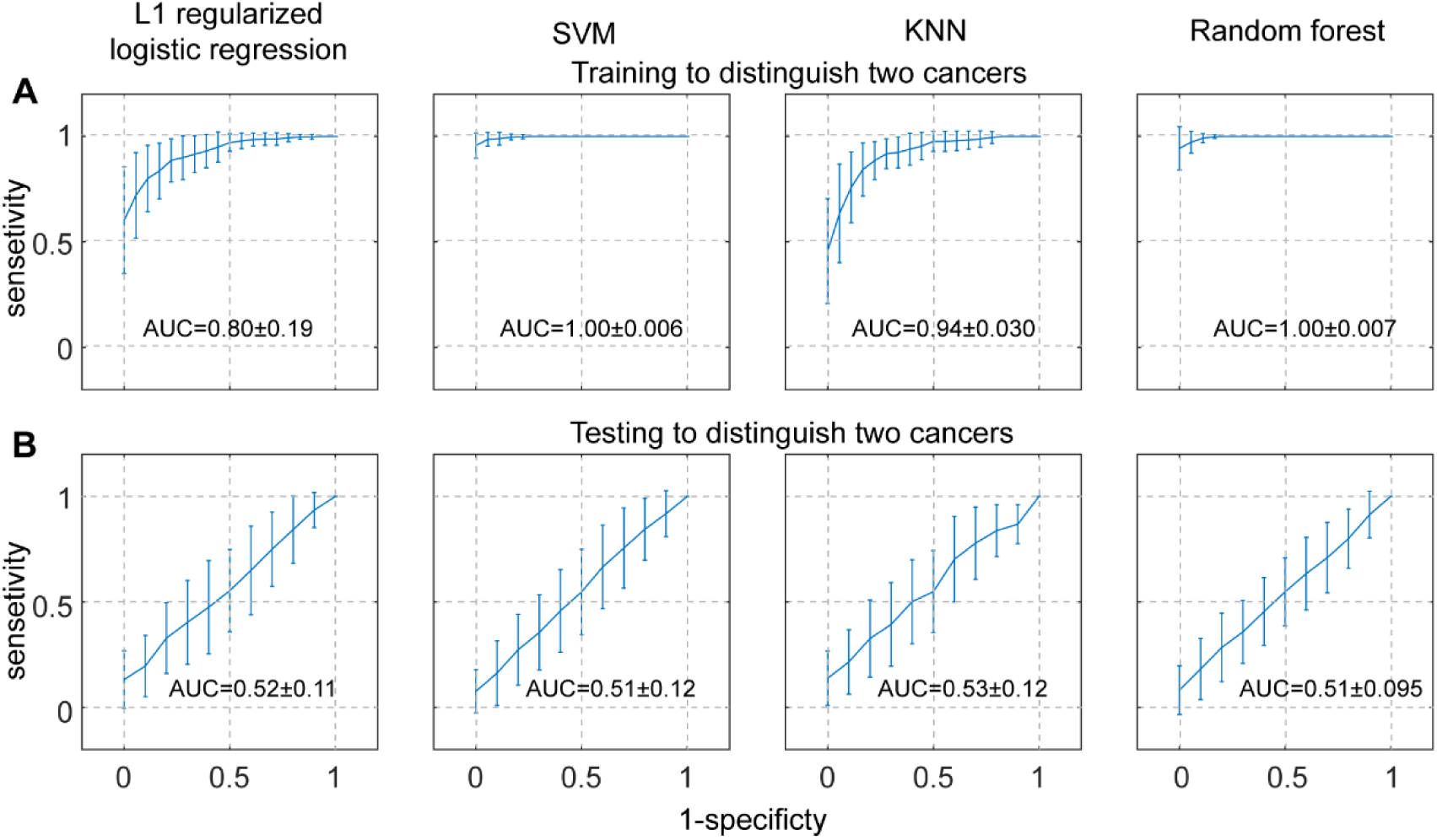
Performances of distinguishing lung cancer and colon cancer plasma RRBS. (A) Training ROC curves for distinguishing lung cancer plasma and colon cancer plasma (RRBS). Standard derivation bars are labeled showing the performance variations of 50 runs. AUCs are labeled on the plots. (B) Testing ROC curves for distinguishing lung cancer plasma and colon cancer plasma (RRBS).

